# Hyperactive BMP and Mechanosignaling Remodel Chromatin to Drive Aberrant Osteogenesis in FOP

**DOI:** 10.64898/2026.01.05.697733

**Authors:** Ellen Y. Zhang, Douglas W. Roberts, Sang Hyun Lee, Bat-Ider Tumenbayar, Jaeun Jung, Loreilys Mejías Rivera, Inkyung Jung, Robert L. Mauck, Eileen M. Shore, Su Chin Heo

## Abstract

Fibrodysplasia ossificans progressiva (FOP) is a rare genetic disorder in which a recurrent ACVR1 (R206H) mutation drives progressive heterotopic ossification (HO). While aberrant BMP hypersensitivity has been studied, how this mutation enforces a persistent pro-osteogenic state remains unclear. Here, we combined super-resolution stochastic optical reconstruction microscopy (STORM), transposase-accessible chromatin with sequencing (ATAC-Seq), and RNA sequencing (RNA-Seq) to investigate how *Acvr1^R206H^* remodels chromatin to promote osteogenic transcriptional programs. Mutant mouse embryonic fibroblasts (MEFs) exhibited globally decondensed chromatin and increased accessibility at developmental and osteogenic loci enriched for HOX, TEAD, and RUNX motifs. Integration of ATAC-Seq and RNA-Seq data identified transcriptional networks primed for osteochondrogenic gene expression, including ossification, extracellular matrix organization, and cell adhesion pathways, consistent with enhanced BMP–SMAD and mechanotransduction activity. Time-course experiments revealed heightened responses to BMP ligands in Acvr1^R206H/+^ MEFs compared to wild-type, highlighting ligand hypersensitivity. Importantly, pharmacological modulation showed that chromatin alterations were dynamic and reversible: activation of Rho/ROCK in wild-type cells reproduced the mutant chromatin state, while inhibition of Rho/ROCK or BMP–SMAD signaling restored condensation to wild-type levels in mutant cells. Together, these findings establish that *Acvr1^R206H^* enforces a pro-osteogenic chromatin landscape through convergent BMP–SMAD and Rho/ROCK signaling, predisposing progenitors to aberrant differentiation trajectories. Our study reframes FOP as a disorder of persistent, but reversible, chromatin states and identifies novel therapeutic opportunities to restore mesenchymal cell homeostasis and prevent pathological bone formation.

**Significance Statement:** Fibrodysplasia ossificans progressiva (FOP) is a rare genetic disorder in which a mutation in ACVR1/ALK2 drives progressive heterotopic ossification. However, how this mutation enforces a persistent pro-osteogenic state is unclear. Here, we show that the *Acvr1^R206H^* mutation remodels chromatin architecture and accessibility through hyperactive BMP–SMAD and Rho/ROCK signaling, activating transcription factor networks that drive osteochondrogenic gene expression. These chromatin changes are dynamic and reversible with targeted pathway inhibition, revealing therapeutic potential to restore mesenchymal cell plasticity and prevent pathological bone formation.

## Introduction

Fibrodysplasia Ossificans Progressiva (FOP) is a rare, severely debilitating genetic disorder characterized by progressive heterotopic ossification (HO), in which bone forms ectopically within soft connective tissues such as skeletal muscle, tendons, and ligaments (1, 2). Hallmark features include congenital malformations of the great toes and episodic flare-ups of HO from early childhood that progressively lead to joint immobilization and profound functional impairment (3–5). The classic clinical form of FOP is caused by heterozygous mutations in *ACVR1/ALK2*, a type I bone morphogenic protein (BMP) receptor. The most common mutation (c.617G>A; R206H) occurs in 97% of patients and localizes to the receptor’s glycine-serine (GS) activation domain (6, 7). Despite extensive research, there are currently no effective treatments to prevent HO, and surgical excision, unfortunately, often exacerbates disease progression. Thus, there is an urgent need for therapies targeting the root molecular drivers of pathological ossification.

Over the past decade, two central pathological mechanisms have emerged in FOP: hyperactive BMP signaling and dysregulated Rho/ROCK-mediated mechanotransduction (8–12). In healthy cells, BMP ligands initiate signaling by binding to heterotetrameric receptor complexes composed of type I (e.g. ALK2) and type II receptors. The constitutively active type II receptor phosphorylates and activates the GS domain of the type I receptor only upon ligand binding, triggering downstream signaling through both canonical SMAD-dependent pathways (SMAD1/5/8-SMAD4) and non-canonical SMAD-independent pathways (MAPK, PI3K, Rho/ROCK) to coordinate skeletal development (13, 14). These pathways collectively regulate transcriptional programs governing cellular differentiation and fate (15–17). In FOP, the R206H mutation in *ACVR1* drives ligand-independent activation, increases sensitivity to BMP ligands, and confers increased responsiveness to Activin A (2, 18). Mutant ACVR1 additionally confers altered ACVR1 GS domain phosphorylation requirements (18). Using an Acvr1^R206H/+^ knock-in mouse model, we previously demonstrated that this gain-of-function mutation alone recapitulates key human FOP features and drives aberrant chondrogenic and osteogenic differentiation in muscle tissue following injury (19).

Beyond BMP signaling, the *Acvr1^R206H^* mutation also perturbs mechanotransduction. We previously demonstrated that Acvr1^R206H/+^ mouse embryonic fibroblast (MEFs) exhibit elevated RhoA/ROCK activity and increased cytoskeletal tension, which were associated with enhanced osteogenic potential in mesenchymal progenitors (11). Notably, Rho/ROCK activation also promoted nuclear translocation of YAP/TAZ, transcriptional co-activators that synergize with BMP–SMAD to amplify osteogenic transcription (11). Together, these findings suggest a reinforcing loop in which BMP hyperactivation and dysregulation of mechanotransduction synergistically exacerbate the propensity of progenitor cells to form HO. However, how these signaling abnormalities remodel chromatin architecture, a key regulator of transcriptional output, remains unknown.

Chromatin organization plays a pivotal role in gene expression, modulating the accessibility of transcriptional machinery to genomic regulatory elements (20). Chromatin architecture and accessibility states are dynamic and responsive to extracellular cues, including both biochemical stimuli (e.g., BMP ligands) and mechanical forces (21, 22). In FOP, chromatin remodeling may reinforce osteogenic lineage commitment in mesenchymal progenitors, sustaining pathogenic gene expression, even after transient signaling events. Thus, understanding how chromatin accessibility and nuclear architecture are altered in Acvr1^R206H/+^ cells is critical for elucidating why these cells are predisposed to aberrant lineage commitment and form pathological bone in soft tissues, which could inform the identification of potential therapeutic targets. Many factors can impact chromatin structure, including mechanical forces transmitted via the cytoskeleton that directly deform the nucleus and alter chromatin compaction (21, 22). Our previous work demonstrated that Acvr1^R206H/+^ MEFs exhibit elevated cytoskeletal tension via Rho/ROCK signaling, suggesting mechanical signaling contributes to chromatin alterations in FOP (11). While both BMP and Rho/ROCK signaling are known to impact chromatin states, their specific and potentially distinct effects in the context of the FOP mutation have not been systematically explored.

Therefore, here, using super-resolution microscopy in combination with RNA sequencing (RNA-Seq) and Assay for Transposase-Accessible Chromatin with Sequencing (ATAC-Seq) analyses, we investigated how BMP stimulation and Rho/ROCK inhibition independently regulate chromatin organization in Acvr1^R206H/+^ cells. Our findings reveal distinct epigenetic mechanisms by which these dysregulated pathways may contribute to pathological ossification. Furthermore, these findings suggest potential therapeutic avenues, as pharmacological targeting of chromatin architecture or mechanotransduction pathways may help restore normal gene expression and cellular identity in Acvr1^R206H/+^ cells.

## Results

### *Acvr1^R206H^* induced nanoscale chromatin remodeling and transcription factor dysregulation

Wild-type (Acvr1^+/+^) and mutant (Acvr1^R206H/+^) mouse embryonic fibroblasts (MEFs) were used to examine how the *Acvr1^R206H^*mutation alters nuclear architecture and transcription factor (TF) accessibility (**Figure 1A**). To visualize global nanoscale chromatin organization at the single-cell level, we employed super-resolution stochastic optical reconstruction microscopy (STORM) by labeling histone H2B in Acvr1^+/+^ and Acvr1^R206H/+^ MEFs. Quantitative analysis revealed that the *Acvr1^R206H^* mutation significantly reduced local H2B Voronoi polygon density, indicating a decrease in chromatin condensation compared to wild-type MEFs (**Figure 1B, C**) and suggesting a more transcriptionally active state. Given the enhanced osteogenic potential of FOP cells, we hypothesized that this chromatin decondensation may reflect a genome primed for lineage-specific differentiation.

**Figure 1.**
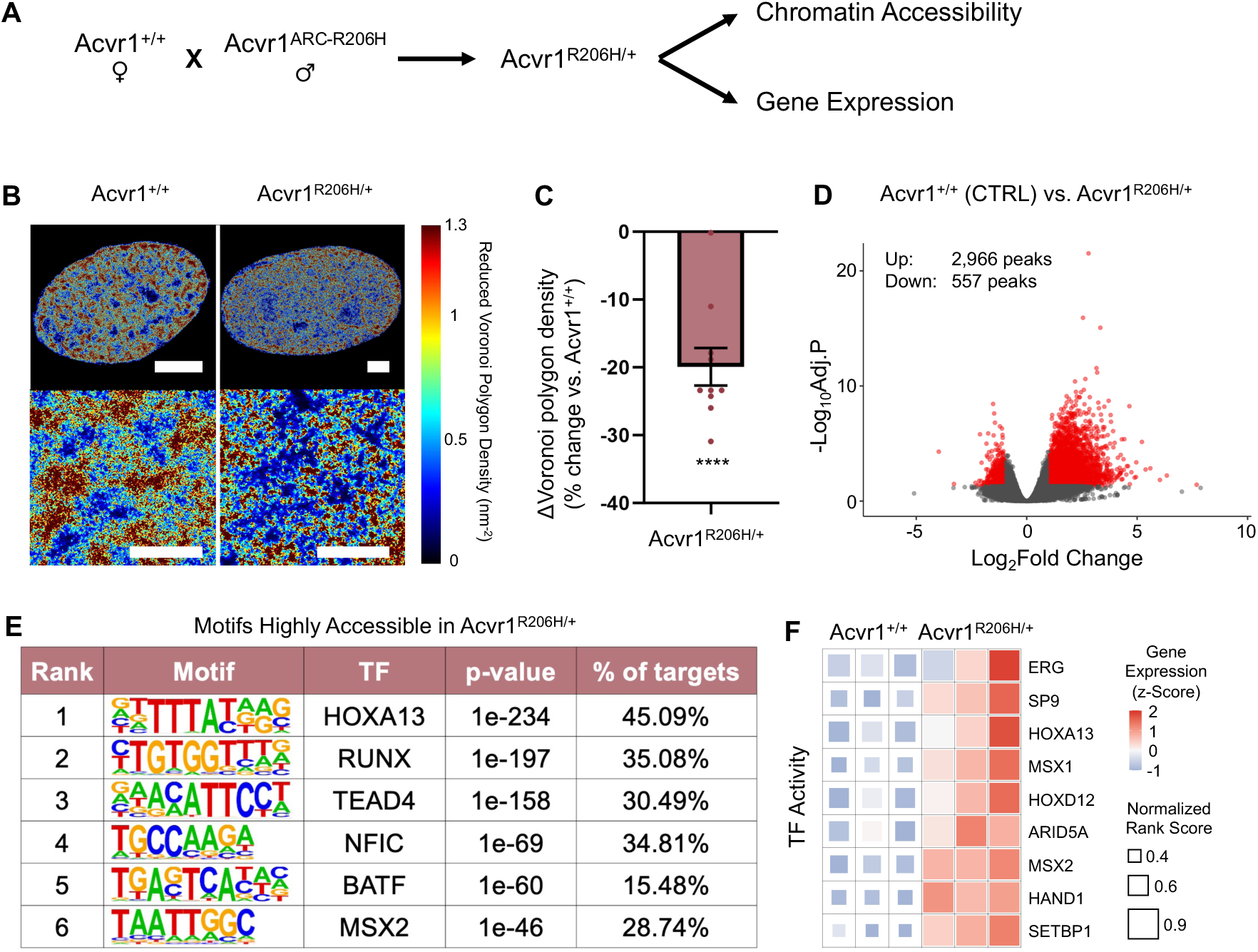
Differences in nanoscale chromatin organization and transcription factor activity between Acvr1^+/+^ and Acvr1^R206H/+^ MEFs. **(A)** Schematic showing breeding of mouse model. **(B)** STORM-based Voronoi polygon density heatmaps of chromatin organization in Acvr1^+/+^ and Acvr1^R206H/+^ MEFs. Scale bars: 5 μm (top), 1 μm (bottom). **(C)** Percentage change in chromatin condensation level, showing decreased chromatin compaction in Acvr1^R206H/+^ MEFs (n = 9) compared to Acvr1^+/+^ MEFs (n = 10). Data indicate mean ± SEM; ****p < 0.0001 (Student’s t-test). **(D)** Volcano plot showing differentially accessible genomic loci in Acvr1^R206H/+^ MEFs, with significantly open and closed regions highlighted in red (n=3 biological replicates). **(E)** De novo transcription factor motif enrichment analysis of genomic loci highly accessible in Acvr1^R206H/+^ MEFs. **(F)** Heatmap showing transcription factor activity at genes that are highly expressed and accessible in Acvr1^R206H/+^ MEFs. Each box represents the z-score of inferred TF activity per condition.

To further assess loci-specific accessibility changes, we performed a bulk assay for transposase-accessible chromatin with sequencing (ATAC-Seq) in Acvr1^+/+^ and Acvr1^R206H/+^ MEFs. Differential accessibility analysis identified 2966 genomic regions with increased accessibility in mutant cells versus only 557 in wild-type cells (**Figure 1D, Supplemental Table S3 and S4**). These results were consistent with our STORM findings that *Acvr1^R206H^* globally decondensed chromatin and enhanced transcriptional potential. To identify potential regulators driving these chromatin accessibility changes, we next performed de novo TF motif enrichment analysis on differentially accessible loci. This motif analysis identified Homeobox A13 (HOXA13), Runt-related transcription factor (RUNX) and TEA domain family member 4 (TEAD4), Nuclear factor I C (NFIC), Basic leucine zipper ATF-like transcription factor (BATF) and MSH homeobox 2 (MSX2) as the top enriched TF motifs in Acvr1^R206H/+^ MEFs (**Figure 1E**). Functionally, HOXA13 plays a crucial role in limb development (23), and its increased accessibility could relate to the congenital great toe malformations characteristic of FOP. In addition, RUNX transcription factors are key regulators of osteogenesis and chondrogenesis (24). The TEAD family of proteins are DNA-binding transcription factors; the mechanosensitive effectors YAP/TAZ serve as co-activators that bind TEAD to drive gene expression (25) associated with osteogenesis (26) and fibrogenesis (27). Finally, we integrated ATAC-Seq and RNA-Seq datasets to infer TF activity changes attributable to *Acvr1^R206H^*. (28). This analysis revealed 9 key TFs, including HOXA13, TEAD4, and multiple RUNX family members, with elevated activity and strong influence over downstream gene expression programs (**Figure 1F**).

Taken together, these data demonstrate that *Acvr1^R206H^* drove global chromatin decondensation, increased accessibility at osteogenic and developmental regulatory loci, and activated a TF network that may underlie both the skeletal malformations and heightened osteogenic potential of Acvr1^R206H/+^ cells.

### *Acvr1^R206H^* enhances BMP–SMAD, cytoskeletal, and chromatin regulatory pathways

Given that *Acvr1^R206H^* alters chromatin organization and increases accessibility at osteogenic and developmental regulatory loci, we next examined whether these changes were associated with transcriptional reprogramming in FOP cells. Bulk RNA sequencing (RNA-Seq) revealed that Acvr1^+/+^ and Acvr1^R206H/+^ MEFs had distinct transcriptomic profiles (**Figure 2A, Supplemental Figure S1A, B, Supplemental Table S1 and S2**). Gene ontology (GO) enrichment analysis identified “ossification”, “extracellular matrix organization”, and “cell adhesion” as the most significantly upregulated biological processes in mutant cells (**Figure 2B, Supplemental Figure S1C**). Given that *Acvr1^R206H/+^* cells exhibited increased response to changes in tissue rigidity (11), we specifically examined genes related to cell adhesion and found that nearly all were significantly upregulated in Acvr1^R206H/+^ MEFs (**Figure 2D**). In addition, Bmp4 was strongly upregulated in Acvr1^R206H/+^ MEFs (**Figure 2E**), along with multiple canonical BMP and WNT target genes, including Msx1, Msx2, Kcp, and Sfrp1/4 (**Figure 2E**). KEGG pathway analysis further confirmed significant enrichment of BMP and WNT signaling pathways in Acvr1^R206H/+^ MEFs (**Figure 2C**). We also observed strong upregulation of extracellular matrix genes associated with chondrogenesis and osteogenesis, including type I collagen (Col1a1, Col1a2), type II collagen (Col2a1), and type VI collagen (Col6a1, Col6a2) in Acvr1^R206H/+^ MEFs (**Figure 2F**). To determine whether these transcriptional changes are linked to chromatin-associated regulatory mechanisms, we compared genes annotated under chromatin remodeling (GO:0006338) with upstream regulators predicted by Ingenuity Pathway Analysis (IPA). This overlap identified 10 key transcriptional modulators within the Acvr1^R206H/+^ gene expression program, including EZH2, CTCF, and GATA3 (**Figure 2G, H**). Interaction network analysis centered on EZH2 further revealed predicted connections to extracellular matrix components (COL1A1, COL1A2, TGFBI), BMP/WNT pathway members (WNT7B, BMP4), and osteogenic regulators (SOX9, ALPL) (**Figure 2I**). Consistent with these transcriptomic predictions, EZH2 (a histone methyltransferase) protein levels were reduced in Acvr1^R206H/+^ MEFs compared to wild-type controls, as confirmed by western blot analysis (**Supplemental Figure S2A, B**). In addition, Acvr1^R206H/+^ MEFs exhibited decreased gene expression related to multiple histone methyltransferases, accompanied by increased expression of histone demethylases (**Supplemental Figure S2C, D**). These results indicate that the Acvr1^R206H^ mutation may disrupt epigenetic homeostasis, leading to altered chromatin organization and accessibility (**Figure 1B-D**), which may contribute to osteogenic reprogramming.

**Figure 2.**
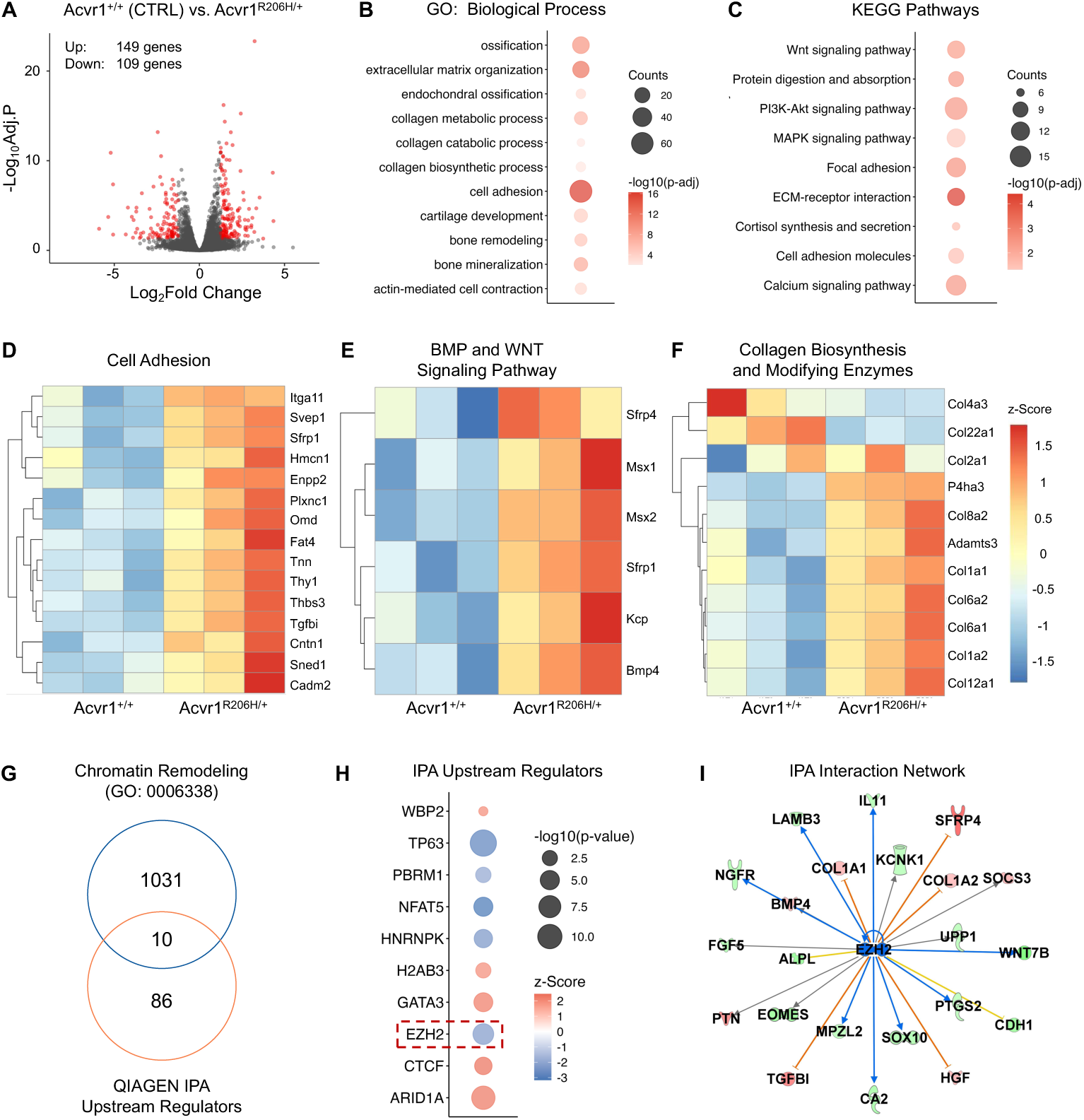
Transcriptomic differences between Acvr1^+/+^ and Acvr1^R206H/+^ MEFs. **(A)** Volcano plot showing differentially expressed genes (DEGs) in Acvr1^R206H/+^ MEFs, with significantly upregulated and downregulated genes highlighted in red (n=3 biological replicates). **(B)** Gene Ontology (GO) enrichment of biological process (BP) and **(C)** KEGG pathway analysis of upregulated genes in Acvr1^R206H/+^MEFs. **(D)** Heatmap of genes related to cell adhesion, **(E)** BMP and WNT signaling pathways, and **(F)** collagen biosynthesis and modifying enzymes in Acvr1^R206H/+^MEFs. **(G)** Venn diagram showing overlap between chromatin remodeling–associated genes (GO:0006338) and predicted upstream regulators identified by Ingenuity Pathway Analysis (IPA). **(H)** IPA upstream regulator analysis highlighting candidate regulators driving transcriptional changes in Acvr1^R206H/+^ MEFs. Dot size represents statistical significance, and color denotes predicted activation z-score. **(I)** IPA-predicted interaction network centered on EZH2, showing connections to extracellular matrix components, BMP/WNT pathway members, and osteogenic regulators. (n=3 biological replicates).

Together, these transcriptomic data indicate that *Acvr1^R206H^* not only amplified BMP–SMAD signaling but also promoted extracellular matrix remodeling and cytoskeletal-related gene expression, thereby altering mechanotransduction pathways. These coordinated changes likely contribute to the heightened osteogenic potential and pathological ossification characteristic of FOP. Moreover, the enrichment and differential expression of chromatin remodeling factors such as EZH2 suggest that epigenetic regulation reinforces these transcriptional programs, linking hyperactive BMP signaling to sustained osteochondrogenic differentiation.

### BMP–SMAD signaling dynamically remodels chromatin structure, with hypersensitive responses in Acvr1^R206H/+^ cells

Given that the *Acvr1^R206H^* mutation hyperactivates BMP signaling activity and previous studies have found that BMP ligand stimulation promotes osteogenic lineage commitment (10), we further investigated how BMP stimulation remodeled chromatin structure in MEFs (**Figure 3A**). To do so, we performed a time-course experiment in which Acvr1^+/+^ and Acvr1^R206H/+^ MEFs were treated with recombinant BMP4 (15 ng/mL), followed by STORM imaging at 0, 0.5, 1, 3, and 12 hours post-treatment (**Figure 3B**). Compared to wild-type cells, Acvr1^R206H/+^ MEFs exhibited a markedly heightened response to BMP stimulation, with significant chromatin decondensation detectable as early as 0.5 hours post-treatment (**Figure 3C, D**). These changes peaked at 3 hours and subsided by 12 hours (**Figure 3C, D**). These results indicate that BMP signaling can rapidly and dynamically remodel chromatin structure, and that Acvr1^R206H/+^ cells were hypersensitive to BMP stimulation, leading to exaggerated and transient chromatin decondensation.

**Figure 3.**
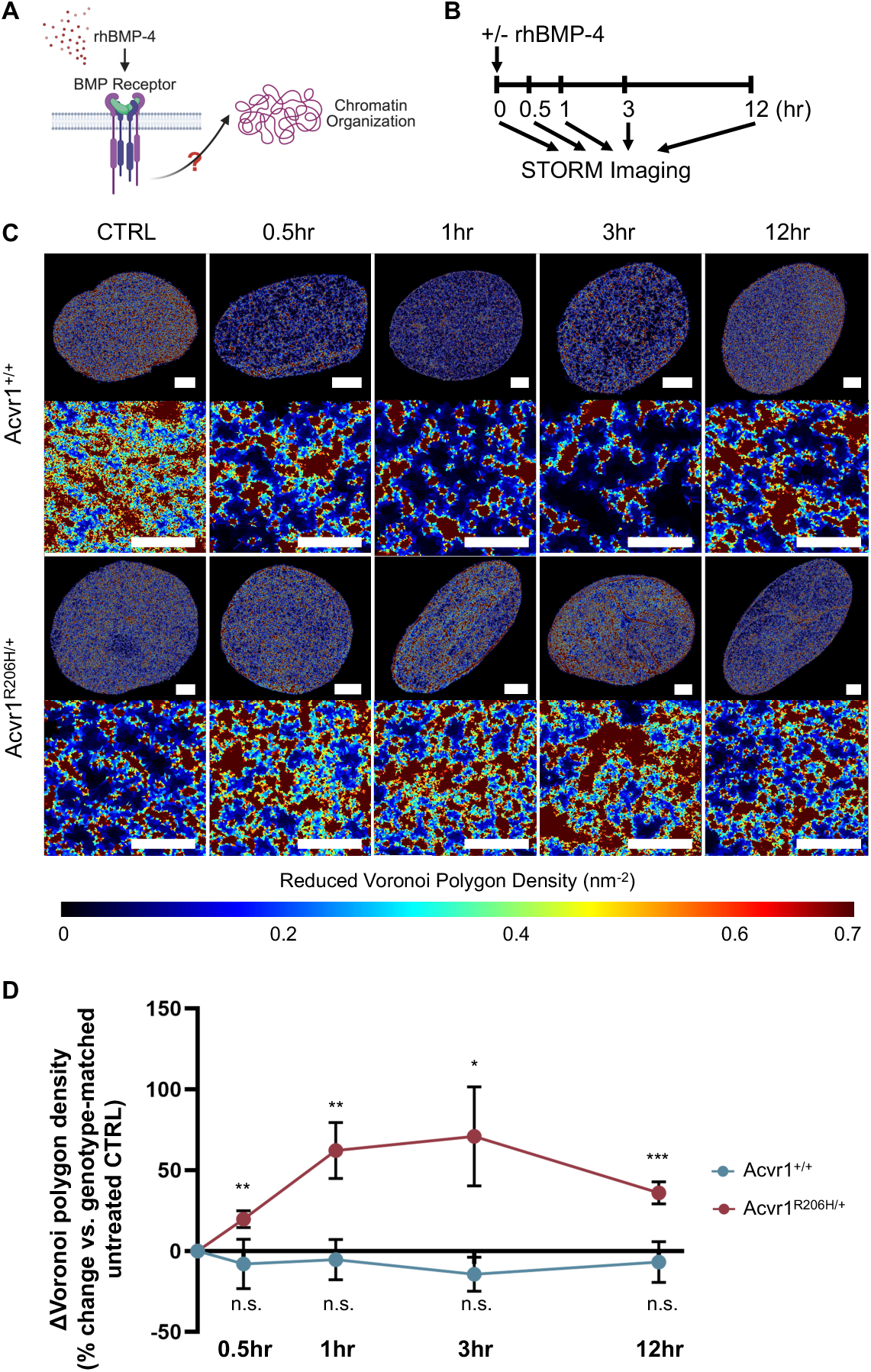
Acvr1^R206H/+^ MEFs are hypersensitive to BMP stimulation. **(A)** Schematic showing study design to test whether BMP stimulation causes chromatin remodeling and **(B)** experimental timeline. **(C)** STORM-based Voronoi polygon density heatmaps of chromatin organization in Acvr1^+/+^ and Acvr1^R206H/+^ MEFs treated with BMP4 at 0, 0.5, 1, 3, and 12 hours post-treatment. Scale bars: 5 μm (top), 1 μm (bottom). **(D)** Percentage change in chromatin condensation level in Acvr1^+/+^ (n=10) and Acvr1^R206H/+^ (n=10) MEFs post-BMP4 treatment, compared to genotype-matched controls. Data indicate mean ± SEM; *p < 0.05, **p < 0.01, ***p < 0.001 (Student’s t-test).

### Pharmacological modulation of BMP**–**SMAD or Rho/ROCK signaling restores chromatin organization

Given that the *Acvr1^R206H^* mutation alters chromatin structure and that both BMP–SMAD and Rho/ROCK pathways are dysregulated in FOP, we next examined the relationship between cytoskeletal tension, BMP signaling, and chromatin organization. To do this, we performed pharmacological interventions targeting each pathway (**Figure 4A, B**).

**Figure 4.**
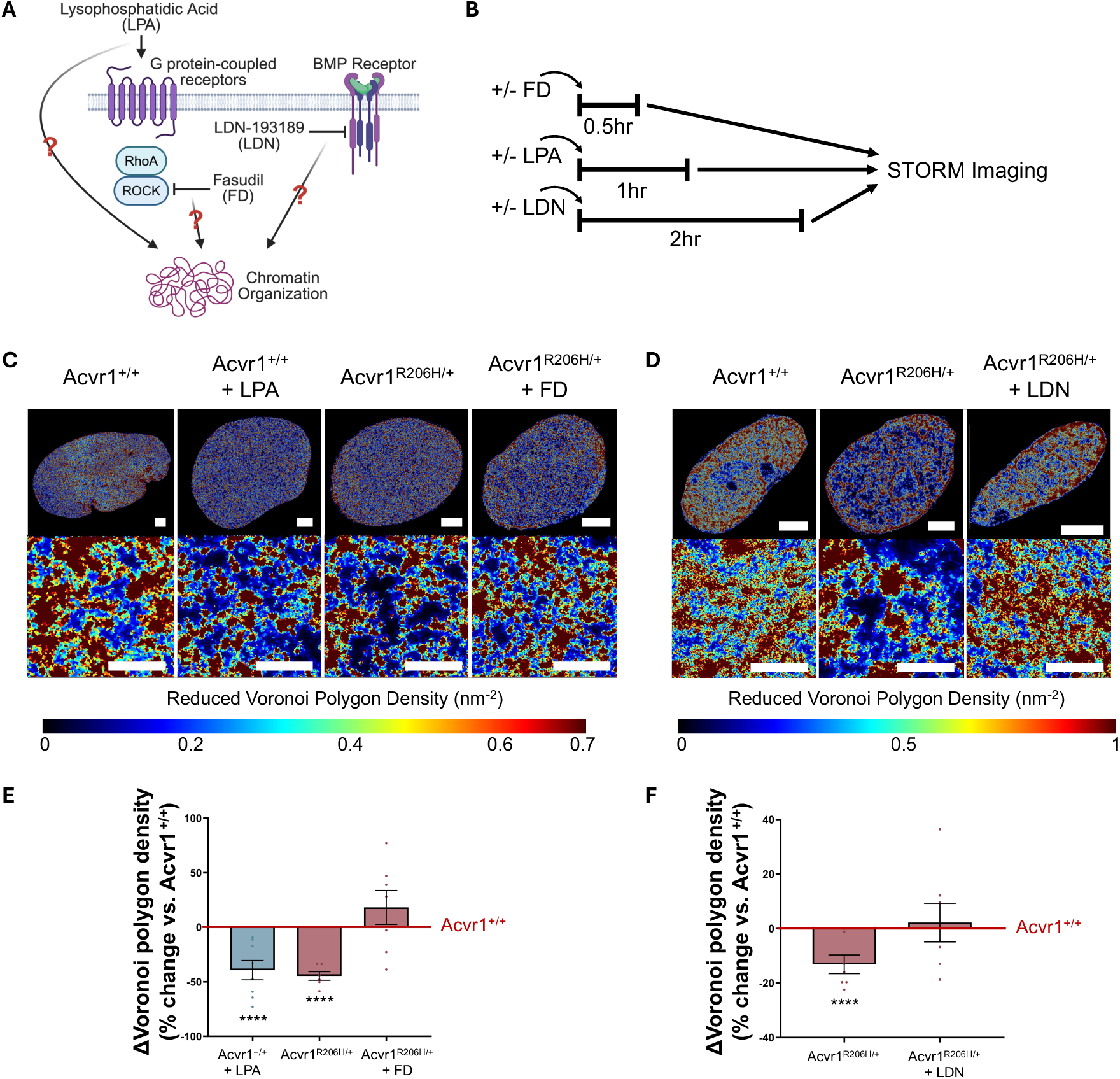
Modulation of BMP–SMAD and Rho/ROCK signaling restores chromatin organization in Acvr1^R206H/+^ MEFs. **(A)** Schematic showing study design to determine if modulation of BMP–SMAD or Rho/ROCK signaling regulates chromatin organization in Acvr1^+/+^ and Acvr1^R206H/+^ MEFs and **(B)** experimental timeline. **(C, D)** STORM-based Voronoi polygon density heatmaps of chromatin organization in Acvr1^+/+^ and Acvr1^R206H/+^ MEFs treated with **(C)** LPA (n=6) or FD (n=7) and **(D)** LDN (n=7). Scale bars: 5 μm (top), 1 μm (bottom). **(E, F)** Percentage change in chromatin condensation level in Acvr1^+/+^ and Acvr1^R206H/+^ MEFs post-LPA/FD **(C)**or LDN **(D)** treatment. Data indicate mean ± SEM; ****p < 0.0001, compared to Acvr1^+/+^ (Student’s t-test).

In wild-type Acvr1^+/+^ MEFs, addition of lysophosphatidic acid (LPA) (which induces Rho/ROCK activation) recapitulated the decondensed chromatin phenotype observed in Acvr1^R206H/+^ MEFs (**Figure 4C, E**). Conversely, treatment of mutant cells with Fasudil (FD), a clinically-used Rho kinase inhibitor, restored chromatin condensation to wild-type levels (**Figure 4C, E**). These results demonstrate that the chromatin structural defects in Acvr1^R206H/+^ MEFs are mechanoresponsive and not irreversible consequences of the mutation. The ability of Rho/ROCK inhibition to normalize both cytoskeletal contractility and chromatin structure suggests this pathway is a critical mechanistic link between the *Acvr1^R206H^* mutation and downstream nuclear architecture alterations.

We next tested whether blocking BMP–SMAD signaling could similarly rescue chromatin organization (**Figure 4A**). Treatment of Acvr1^R206H/+^ MEFs with LDN-193189 (LDN), an inhibitor of SMAD1/5/8 phosphorylation downstream of BMP stimulation (29), restored chromatin condensation to wild-type levels (**Figure 4D, F**). Together, these findings identify Rho/ROCK and BMP–SMAD pathways as key regulators of chromatin structure in Acvr1^R206H/+^ cells and highlight both as promising therapeutic targets for restoring normal nuclear architecture in Acvr1^R206H/+^ MEFs.

## Discussion

Heterotopic ossification (HO) in FOP raises a central question: how does the constitutively active *Acvr1^R206H^* mutation lead to persistent and pathological shifts in mesenchymal cell fate? Although aberrant BMP hypersensitivity signaling has been well documented, the mechanisms by which this mutation establishes and sustains a pro-osteogenic state remain poorly understood, particularly in terms of chromatin architecture and transcriptional regulation.

To address this, here we combined nanoscale super-resolution imaging, epigenomic, and transcriptomic profiling to demonstrate that the *Acvr1^R206H^* mutation induces extensive remodeling of chromatin structure, accessibility, and gene expression. Under basal conditions, mutant MEFs exhibited globally decondensed chromatin and increased accessibility at the developmental and osteogenic regulatory loci enriched for HOX, TEAD, and RUNX motifs. These chromatin changes occurred in the absence of exogenous stimulation, in contrast to wild-type cells which required BMP4 treatment or RhoA activation to produce similar chromatin patterns. Integration of ATAC-Seq and RNA-Seq revealed that these accessible loci were transcriptionally active and enriched for osteochondrogenic regulators, including Runx2, Col1a1, and Col2a1. Consistent with these results, Acvr1^R206H/+^ cells showed downregulation of histone methyltransferases such as EZH2 and upregulation of histone demethylases such as Kdm6a, indicating a global epigenetic shift toward a more transcriptionally permissive chromatin state. This imbalance between methylation and demethylation suggests that the Acvr1^R206H/+^ mutation disrupts the normal maintenance of histone modifications critical for chromatin stability and lineage commitment. Consequently, altered chromatin organization and increased genomic accessibility (**Figure 1B–D**) may prime cells for aberrant activation of osteogenic gene networks.

Together, these findings support a model in which the *Acvr1^R206H^* mutation initiates and reinforces a pro-osteogenic chromatin and transcriptional landscape through cooperative hyperactivation of BMP–SMAD signaling and RhoA/ROCK-driven YAP/TAZ nuclear localization, which cooperate with chromatin-modifying enzymes to activate osteochondrogenic gene expression (**Figure 5**). In the wild-type state, low SMAD nuclear translocation, cytoplasmic sequestration of YAP/TAZ, and repressive chromatin marks deposited by methyltransferases such as EZH2 and SMYD family members (20, 30) may maintain suppression of osteochondrogenic gene expression. In FOP, reduced regulatory constraints on mutant ACVR1, which result in increased phosphorylation of SMAD1/5/8 drives persistent nuclear accumulation (18, 31), while elevated RhoA/ROCK signaling promotes nuclear translocation of YAP/TAZ (32). These transcriptional co-activators, together with RUNX and other osteogenic TFs, engage chromatin-modifying enzymes, likely including histone demethylases to sustain an open, transcriptionally permissive chromatin state that enhances osteochondrogenesis (**Figure 5**).

**Figure 5.**
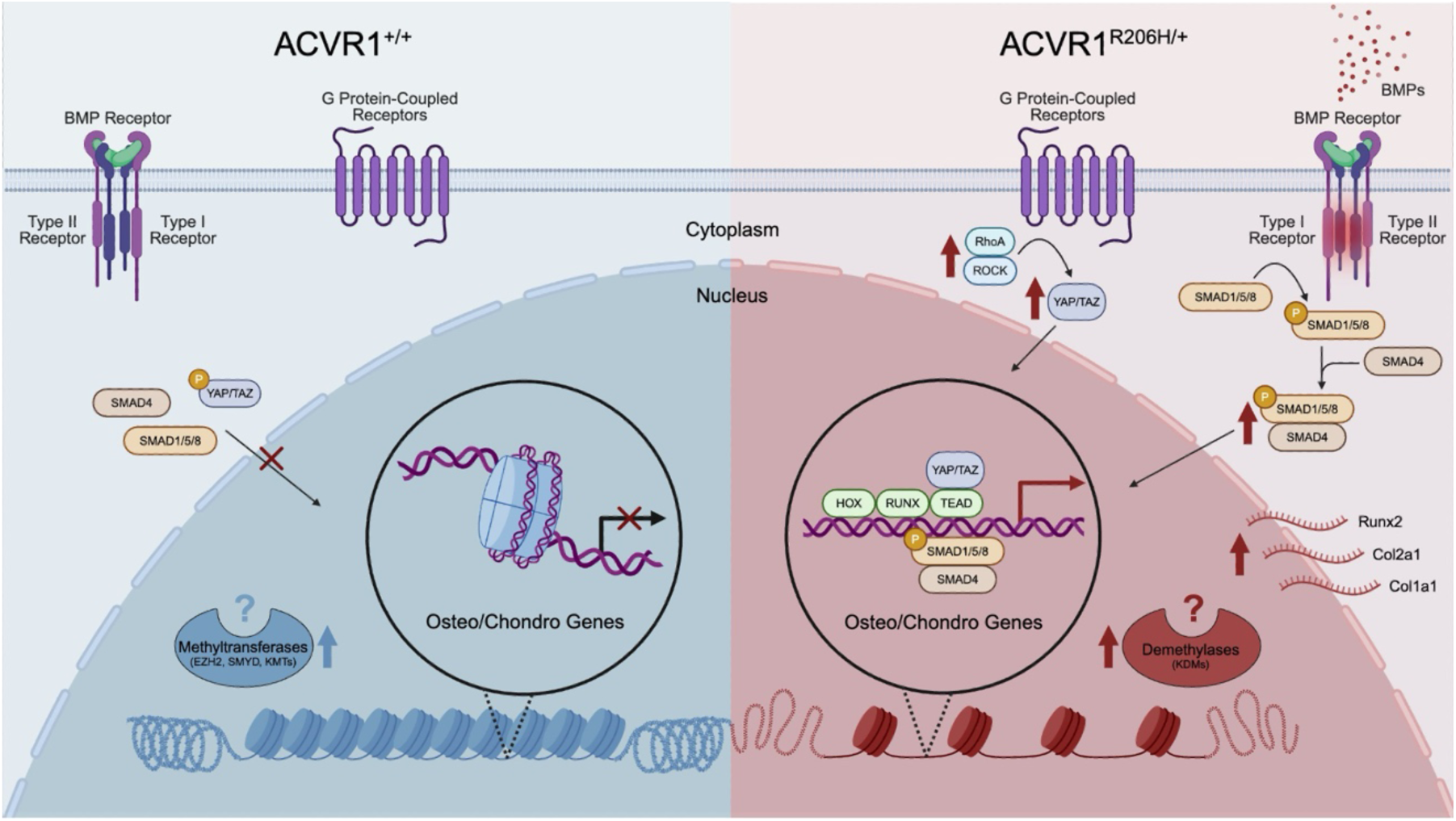
Model of *Acvr1^R206H^*-driven chromatin remodeling and aberrant gene activation in FOP. In wild-type cells (left), BMP–SMAD signaling and YAP/TAZ activity are restrained, and repressive chromatin modifiers maintain osteo/chondro genes in a condensed state. In Acvr1^R206H/+^ cells (right), hyperactive BMP–SMAD signaling and RhoA/ROCK-driven YAP/TAZ nuclear localization cooperate with osteogenic TFs and demethylases to decondense chromatin and activate osteo/chondro genes, driving pathological HO.

Importantly, our pharmacological experiments reveal that these chromatin defects are not fixed consequences of the mutation, but rather are highly mechanoresponsive and reversible. Activation of Rho/ROCK signaling in wild-type cells phenocopied the decondensed chromatin of mutant cells, while inhibition with Fasudil restored chromatin condensation in Acvr1^R206H/+^ MEFs to wild-type levels. Likewise, inhibition of BMP–SMAD signaling with LDN normalized chromatin structure. These results indicate BMP–SMAD and Rho/ROCK pathways are convergent regulators of nuclear architecture in FOP and identify both as promising therapeutic targets.

While our findings define a molecular framework linking constitutive ACVR1 signaling to chromatin remodeling and osteochondrogenic gene activation, several limitations should be acknowledged. Chromatin is a highly dynamic structure, and even in the context of a disease-driving mutation, it retains the potential to reorganize in response to environmental cues (27, 33). For instance, as shown in our previous study (33), disease cells exhibited markedly reduced chromatin dynamics compared to normal cells, suggesting that pathological signaling may constrain nuclear plasticity. In addition, because our experiments used MEFs, which are developmentally plastic, it is possible that cells harboring the ACVR1 mutation for longer periods, may exhibit reduced chromatin flexibility and diminished potential for rescue. Thus, future studies should explore how developmental context influences chromatin remodeling and reversibility of pathogenic signaling. In this study, we observed that chromatin structure was highly responsive to modulations of the BMP–SMAD and Rho/ROCK pathways; however, all experiments were conducted on stiff tissue culture plastic surfaces. This does not replicate the mechanical environment of soft connective tissues where FOP lesions form, and it remains unclear how substrate stiffness may modulate signaling-chromatin interactions *in vivo*. Furthermore, the relationship between BMP–SMAD and Rho/ROCK in the context of the *Acvr1^R206H^* mutation remains incompletely understood. It is not yet known whether these pathways act independently, converge on shared downstream targets, or function in a hierarchical manner to regulate chromatin structure in FOP. Future studies should dissect their interplay, particularly through sequential or combinatorial inhibition experiments in more physiologically relevant culture systems.

Collectively, our study reframes FOP as a disorder of stabilized, yet reversible, bias toward an aberrant fate, following from heightened BMP signaling that converges with epigenetic mechanisms to maintain pro-osteogenic gene expression programs. This perspective suggests new therapeutic opportunities: beyond inhibiting upstream signaling, targeting chromatin modifiers or mechanosensitive transcriptional regulators may reduce osteogenic bias and prevent heterotopic ossification. More broadly, our findings illustrate how persistent signaling dysregulation can reshape the chromatin landscape to drive long-term pathological outcomes, even in the absence of continued extrinsic stimuli.

## Materials and Methods

### Cell Isolation and Culture

Wild-type (Acvr1^+/+^) and mutant (Acvr1^R206H/+^) mouse embryonic fibroblasts (MEFs) were isolated from E13.5 embryos (19, 34). For breeding, control females were crossed with previously recombined heterozygous Acvr1^ARC-R206H^ males. Embryos were dissected, with the head and visceral organs removed, then minced and digested in 0.05% Trypsin-EDTA (Invitrogen) at 37°C for 35 minutes. The resulting cell suspension was transferred to T-75 flasks and cultured in basal growth medium [Dulbecco’s Modified Eagle Medium (DMEM; ThermoFisher, 11965118), supplemented with 10% fetal bovine serum (FBS; R&D Systems, S11150) and 1% penicillin-streptomycin (PS; Corning, 30-002-CI)] at 37°C in a humidified incubator with 5% CO_2_, with media changes every 2–3 days. All experiments were conducted with early-passage cells (passage 1–2).

### Bulk RNA Sequencing (RNA-Seq) and Transcriptome Analysis

Passage 0 Acvr1^+/+^ and Acvr1^R206H/+^ MEFs were cultured for 2 days in six-well plates before RNA extraction using TRIzol Reagent (Invitrogen) and purification with the Direct-zol RNA Microprep Kit (Zymo Research). RNA quality (RIN > 8.0) and concentration were assessed using a NanoDrop spectrophotometer, a Qubit, and a Bioanalyzer (Agilent). Libraries were prepared via rRNA depletion and sequenced as 150 bp paired-end reads on an Illumina platform (Genewiz). Raw reads were trimmed (Trimmomatic) (35) and aligned to mm10 (STAR) (36). Gene counts were generated using featureCounts (Subread) (37). Differential expression analysis (DESeq2) identified differentially expressed genes (DEGs; adjusted p-value < 0.05, |log_2_FoldChange| > 1.0) (38). Results were visualized via EnhancedVolcano and pheatmap (39). Gene Ontology (GO) enrichment (clusterProfiler, FDR ≤ 0.05) was plotted using ggplot2 (40).

### Assay for Transposase-Accessible Chromatin with Sequencing (ATAC-Seq) and Analysis

Nuclei from 1 x 10^5^ Acvr1^+/+^ or Acvr1^R206H/+^ MEFs were isolated (ATAC Lysis Buffer) and tagged using the Active Motif ATAC-Seq Kit (37°C, 30 min). Purified DNA was PCR-amplified (Q5 High-Fidelity Polymerase, NEB) with 10 cycles of indexing, followed by SPRI bead size selection. Library quality was verified via Bioanalyzer (Agilent) and quantified (NEBNext Library Quant Kit). Libraries were prepared and sequenced as 150 bp paired-end reads on an Illumina platform (Genewiz). Raw reads were trimmed (Trimmomatic) (35), aligned to mm10 (Bowtie2) (41), and filtered (SAMtools) (42). Peaks were called (MACS2) (43), merged (bedtools) (44), and quantified (featureCounts) (37).

Differential accessibility analysis (DESeq2) identified differentially accessible peaks (DAPs; adjusted p-value < 0.01, |log_2_FoldChange| > 0) (38). Motif enrichment (HOMER) (45) and key transcription factor (TF) identification (Taiji) (28) were performed, with results visualized with ComplexHeatmap in R.

### Pharmacological Modulation of BMP Signaling and RhoA Activity

To activate BMP–SMAD signaling, Acvr1^+/+^ and Acvr1^R206H/+^ MEFs were treated with 15 ng/mL recombinant human BMP-4 (rhBMP-4; R&D Systems, #314-BP) either overnight or for 0.5–12 h for time-course experiments. BMP signaling was inhibited using LDN-193189 (LDN; 50 nM, 2 h; Sigma, #SML0559) prior to imaging. For RhoA modulation, cells were treated with either the inhibitor Fasudil (FD; 10 µM, 30 min; Sigma, #CDS021620) or the activator lysophosphatidic acid (LPA; 50 µM, 1 h; Sigma, #L7260) prior to imaging.

### Stochastic Optical Reconstruction Microscopy (STORM)

Acvr1^+/+^ and Acvr1^R206H/+^ MEFs were seeded into chambered cover glass and fixed with methanol-ethanol (1:1, -20°C, 6 min), followed by blocking (BlockAid; 1 h, RT; ThermoFisher, #B10710). Chromatin was labeled with anti-H2B antibody (1:50, 4°C, overnight; Invitrogen, #PA5-115361) and Alexa Fluor 405/647-conjugated secondary antibodies (1 h, RT). Imaging was conducted on an ONI Nanoimager using activation-imaging cycles (1 x 405 nm: 3 x 647 nm) in oxygen scavenging buffer (10 mM MEA, 0.5 mg/mL glucose oxidase, 40 mg/mL catalase, 10% glucose) (33, 46–48). Chromatin condensation was quantified via Voronoi tessellation of fluorophore localizations, with smaller polygons indicating higher density (33, 46–48). Nuclear areas were calculated from polygon areas, excluding peripheral regions. For comparative analysis, polygon areas were normalized to their mean and plotted as cumulative distributions. Chromatin domains were identified by clustering adjacent polygons below a size threshold (33, 46–48). All analyses were performed in MATLAB (33, 46–48).

### Statistical analysis

All statistical analyses were performed using GraphPad Prism 10 (La Jolla, CA) and R (38). Data are presented as mean ± SEM with biological replicates (n) indicated in figure legends. For comparisons between two groups, unpaired Student’s t-tests were used. For comparisons among multiple groups, one-way ANOVA with Tukey’s post-hoc test was applied (p-value < 0.05 considered significant).

## Acknowledgments

This research was supported by grants from the National Institutes of Health (AR071399, AR079224) and the NSF Science and Technology Center for Engineering Mechanobiology (CMMI-1578571). Schematic diagrams were created in BioRender (https://biorender.com/e0ke1av)

## Author Contributions

E.Y.Z., D.W.R., S.H.L., B.T., J.J., L.M.R., I.J., R.L.M., E.M.S., and S.C.H. designed research; E.Y.Z., D.W.R., B.T., and J.J. performed research; E.Y.Z., D.W.R., S.H.L., and B.T., contributed new reagents/analytic tools; E.Y.Z., S.H.L., B.T., and J.J. analyzed data; and E.Y.Z., D.W.R., S.H.L., B.T., J.J., L.M.R., I.J., R.L.M., E.M.S., and S.C.H. wrote the paper.

## Competing Interest Statement

The authors declare no competing interest.

## Supporting Information

**Figure S1.**
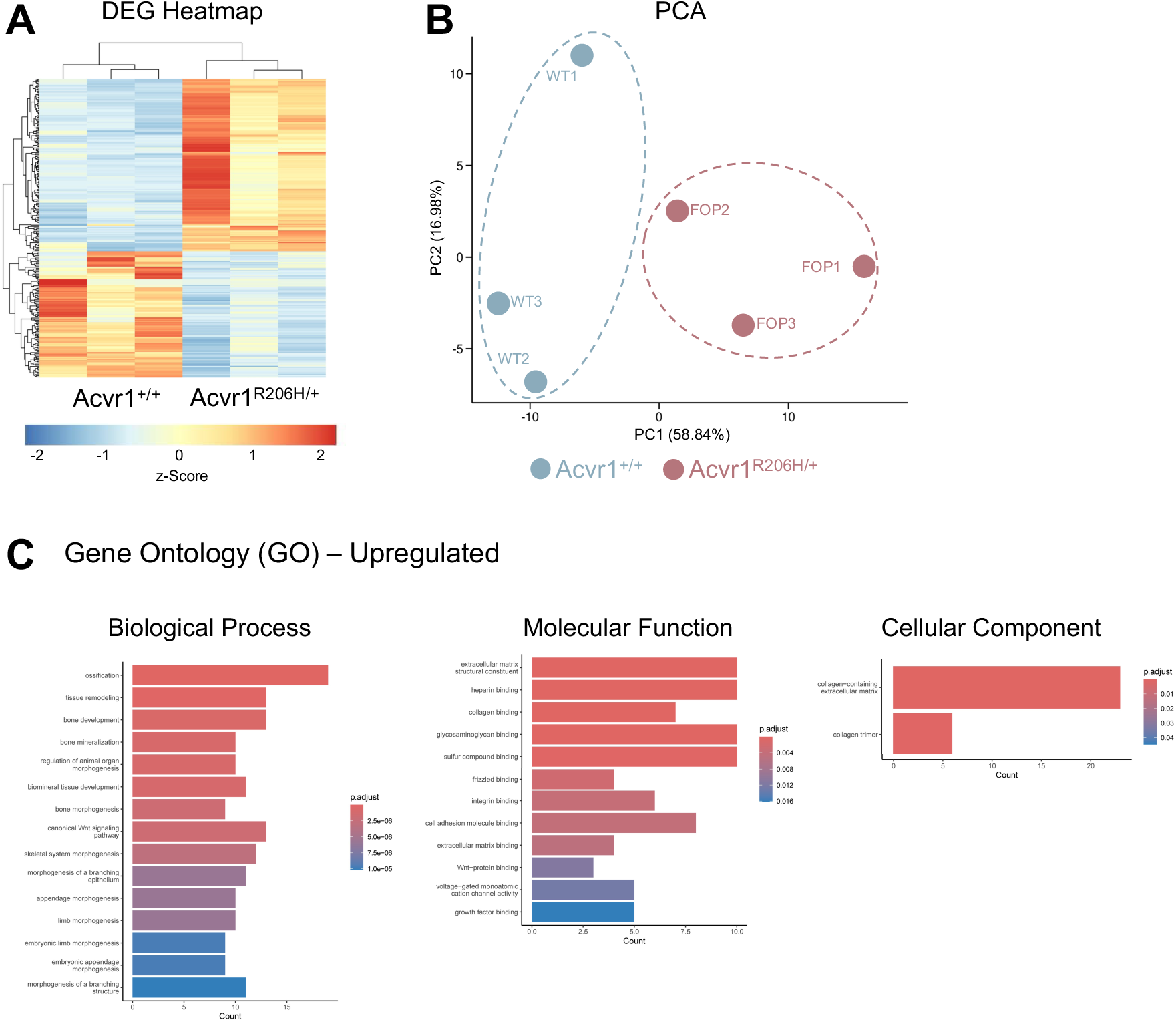
Acvr1^R206H/+^ MEFs display distinct global transcriptomic signatures enriched for osteogenic and matrix-associated pathways. **(A)** Heatmap of differentially expressed genes between Acvr1^+/+^ and Acvr1^R206H/+^ MEFs, showing hierarchical clustering of biological replicates (n = 3 biological replicates). **(B)** Principal component analysis (PCA) plot illustrating distinctions between Acvr1^+/+^ and Acvr1^R206H/+^ MEFs based on global gene expression profiles. **(C)** Gene Ontology (GO) enrichment analysis of upregulated genes in Acvr1^R206H/+^ MEFs, highlighting significantly enriched Biological Process, Molecular Function, and Cellular Component terms.

**Figure S2.**
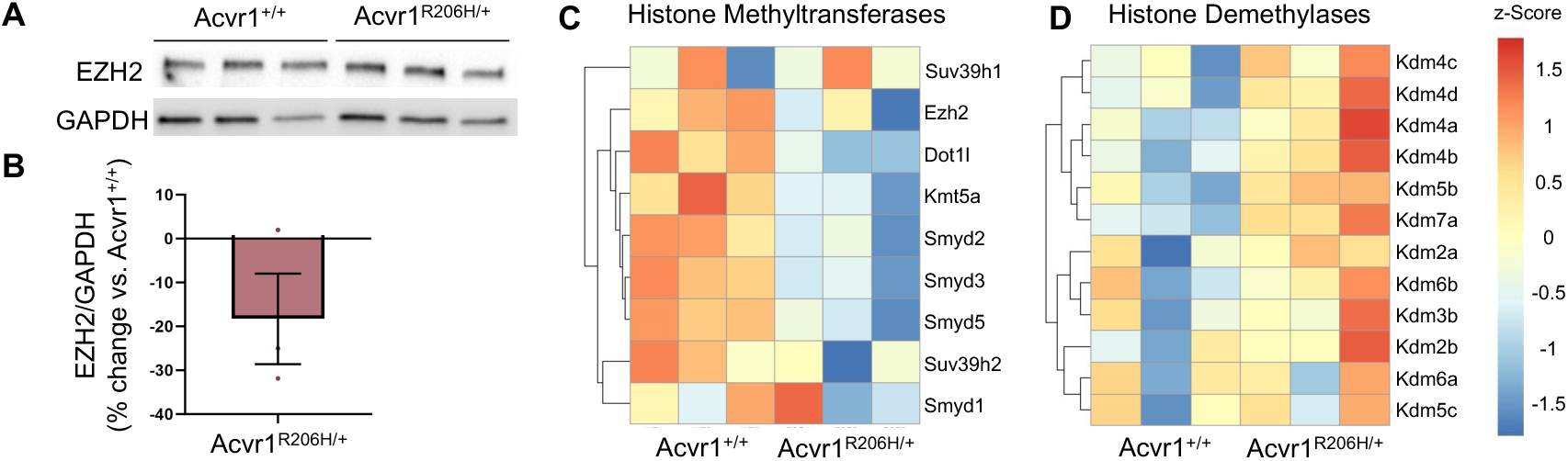
Altered expression of chromatin-modifying enzymes in Acvr1^R206H/+^ MEFs. **(A)** Western blot showing decreased EZH2 protein levels in Acvr1^R206H/+^ MEFs compared to wild-type controls. **(B)** Quantification of relative EZH2 protein expression normalized to GAPDH (n=3 biological replicates). **(C)** Heatmap showing decreased expression of histone methyltransferase genes in Acvr1^R206H/+^ MEFs. **(D)** Heatmap showing increased expression of histone demethylase genes in Acvr1^R206H/+^ MEFs (n=3 biological replicates).

**Table S1.**
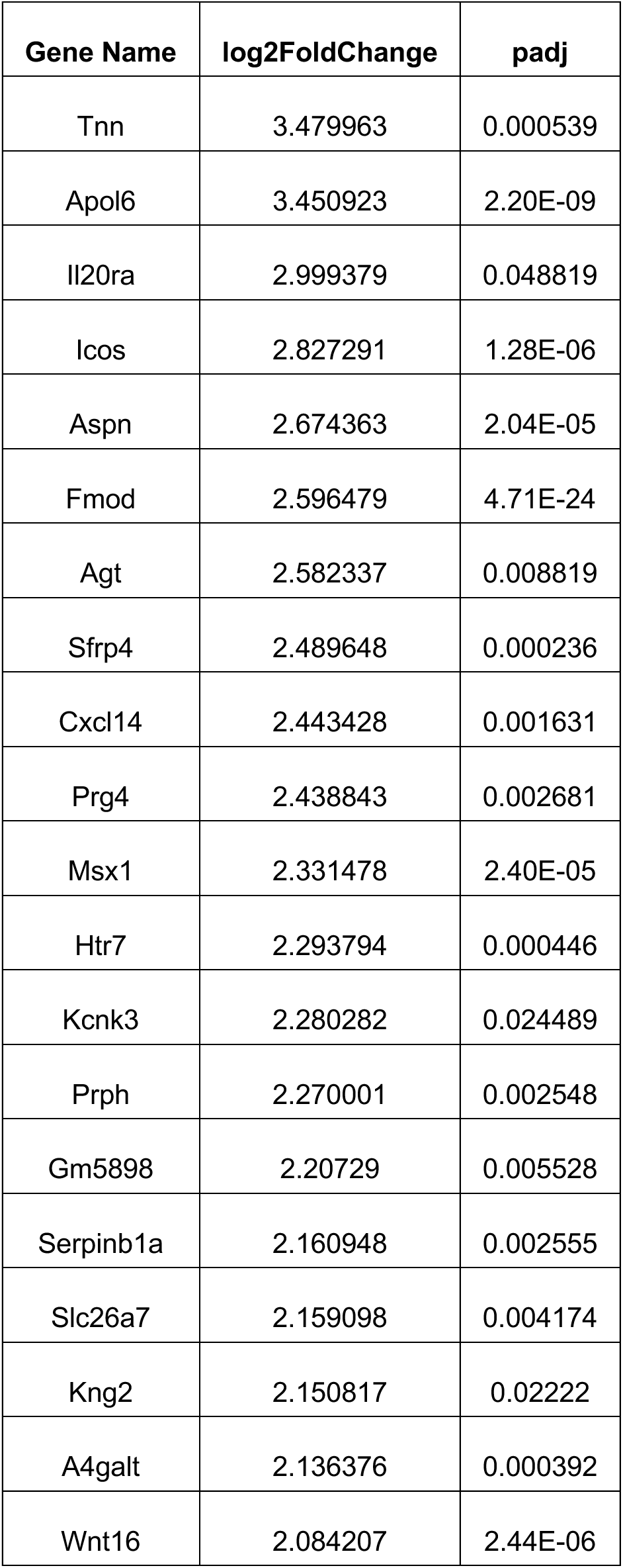
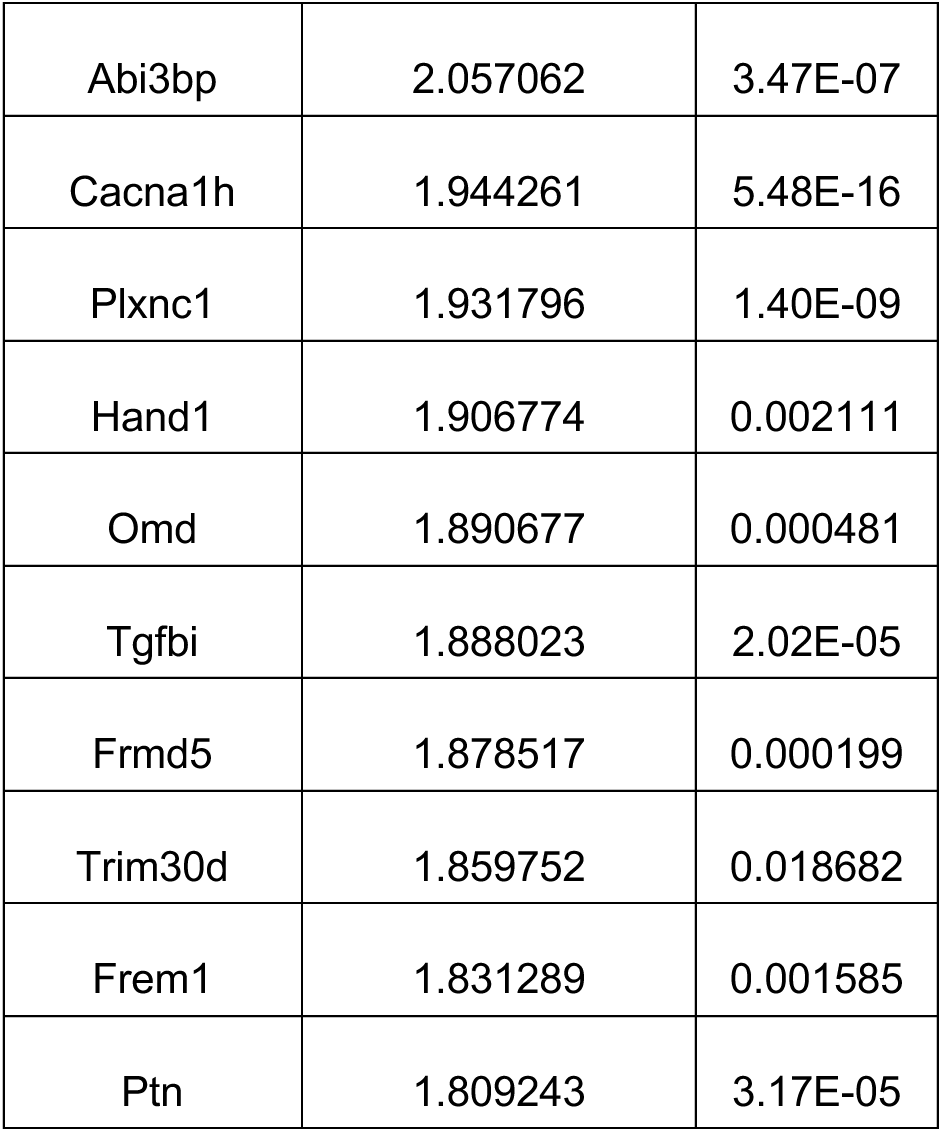
List of top 30 upregulated genes in Acvr1^R206H/+^ MEFs compared to Acvr1^+/+^ MEFs.

**Table S2.**
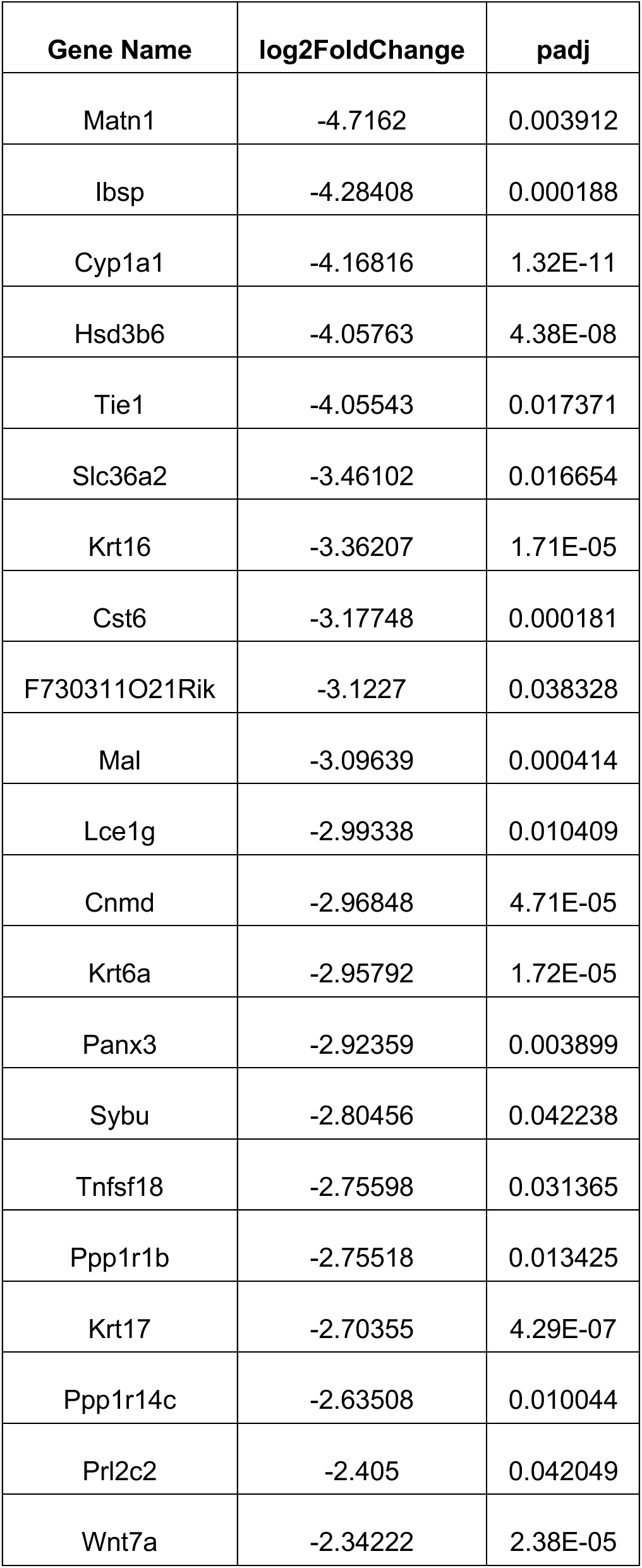
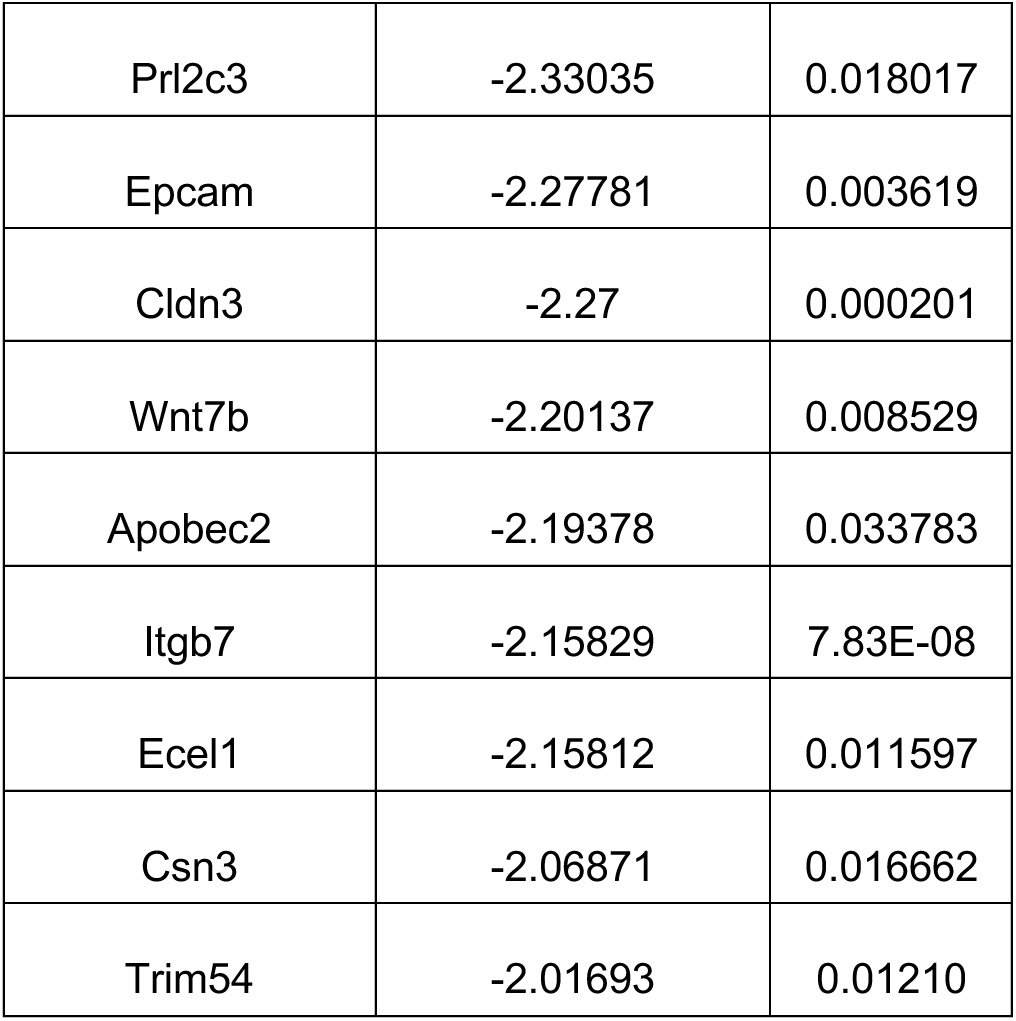
List of top 30 downregulated genes in Acvr1^R206H/+^ MEFs compared to Acvr1^+/+^ MEFs.

**Table S3.**
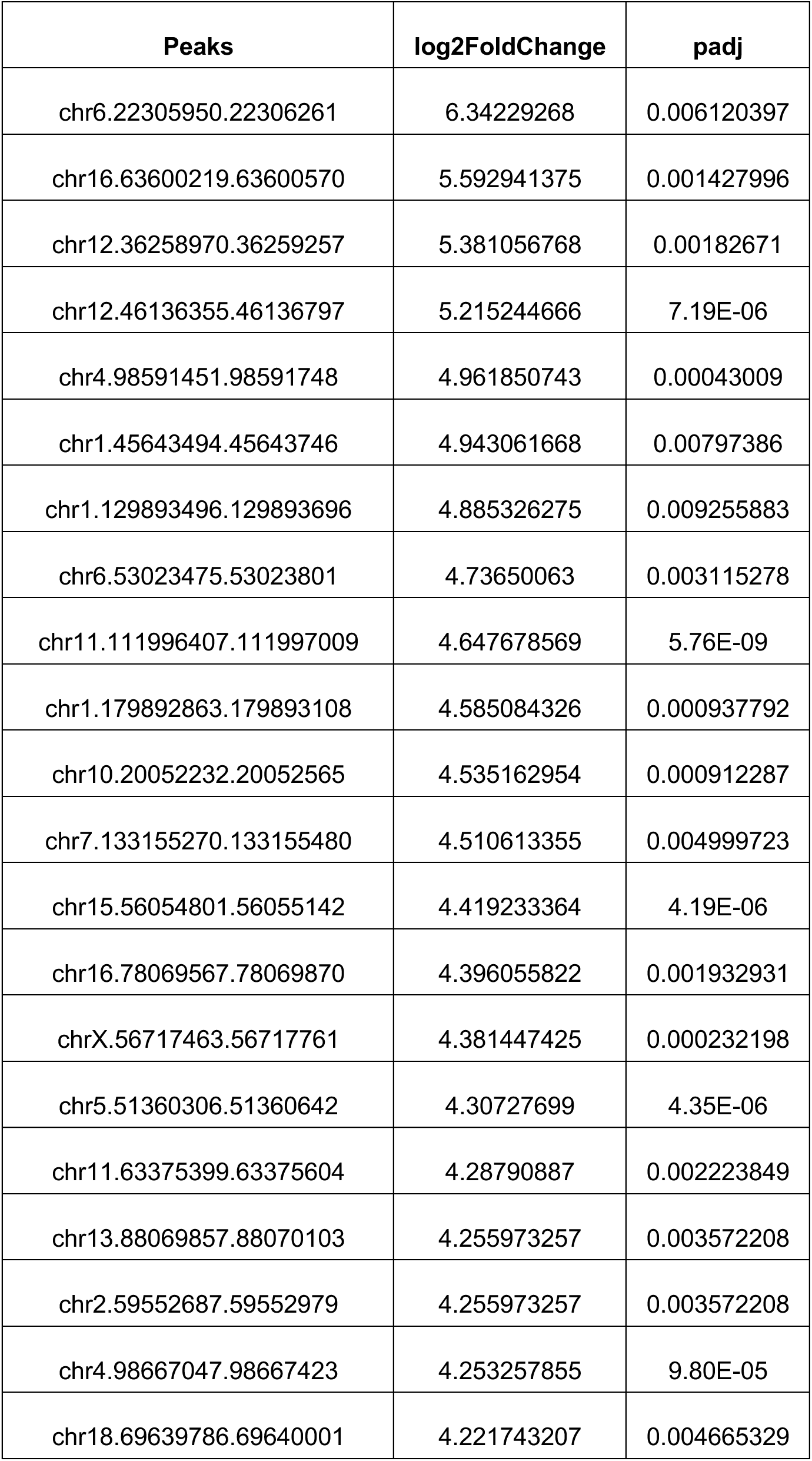
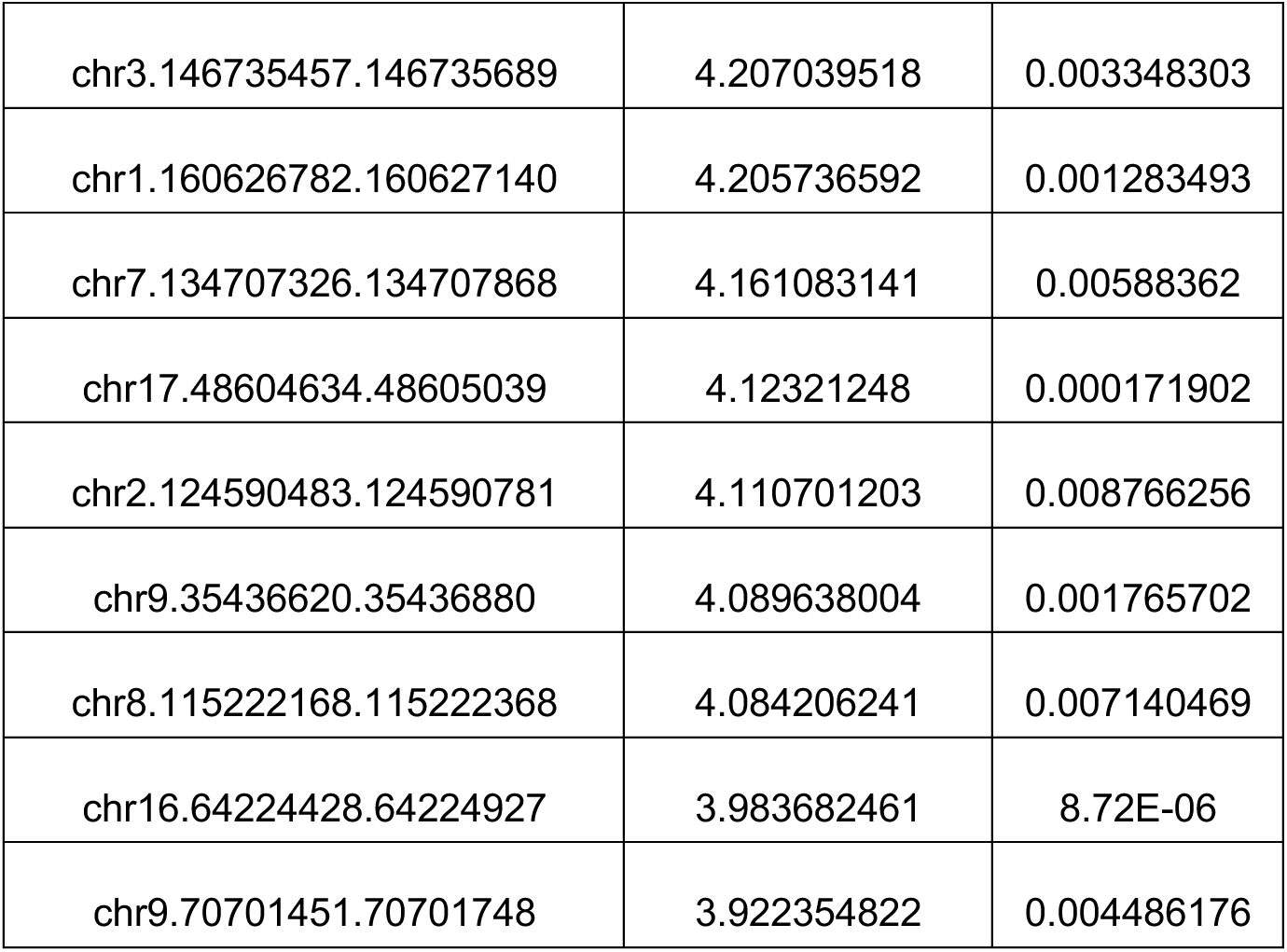
List of top 30 accessible peaks in Acvr1^R206H/+^ MEFs compared to Acvr1^+/+^ MEFs.

**Table S4.**
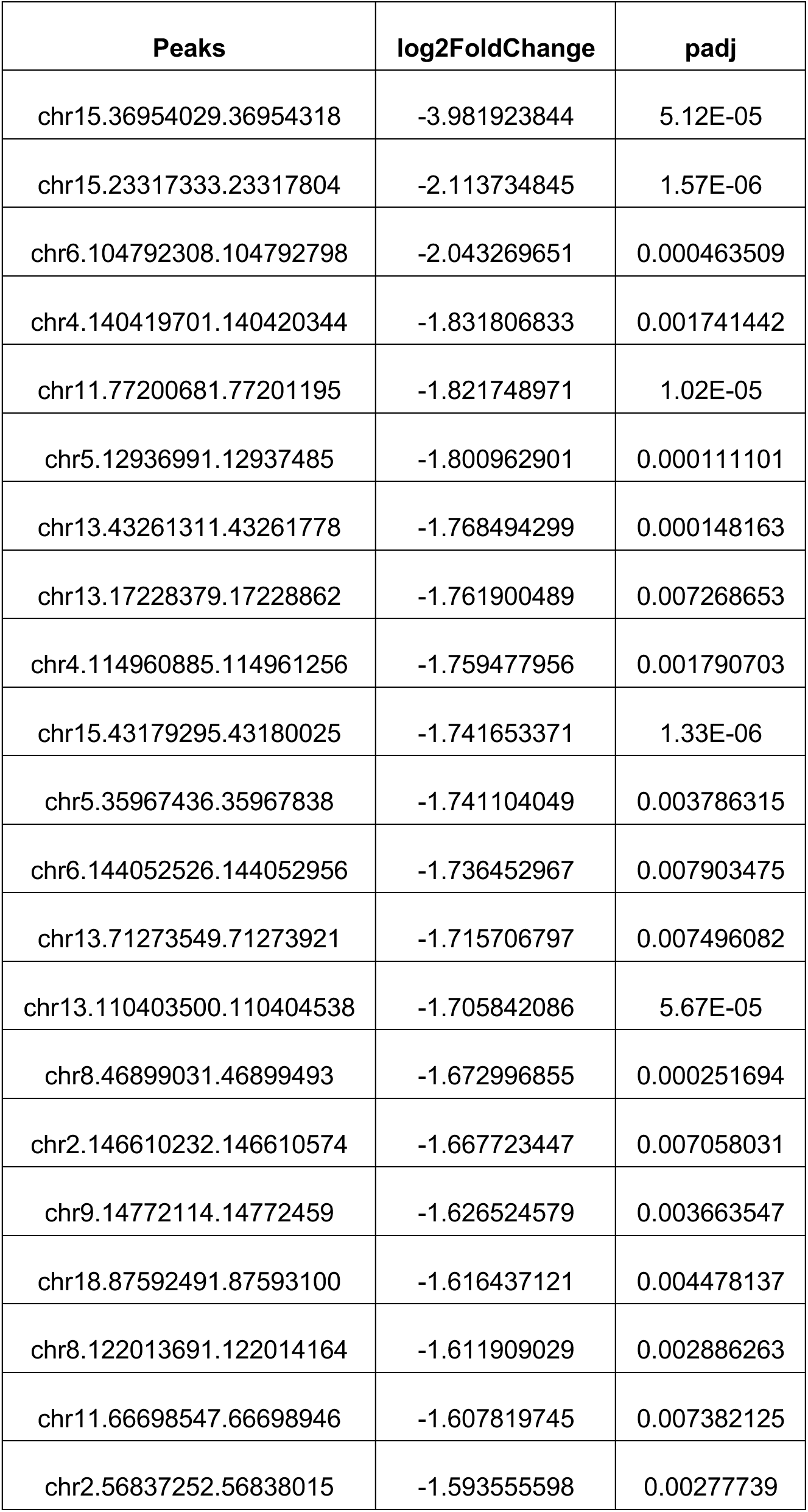
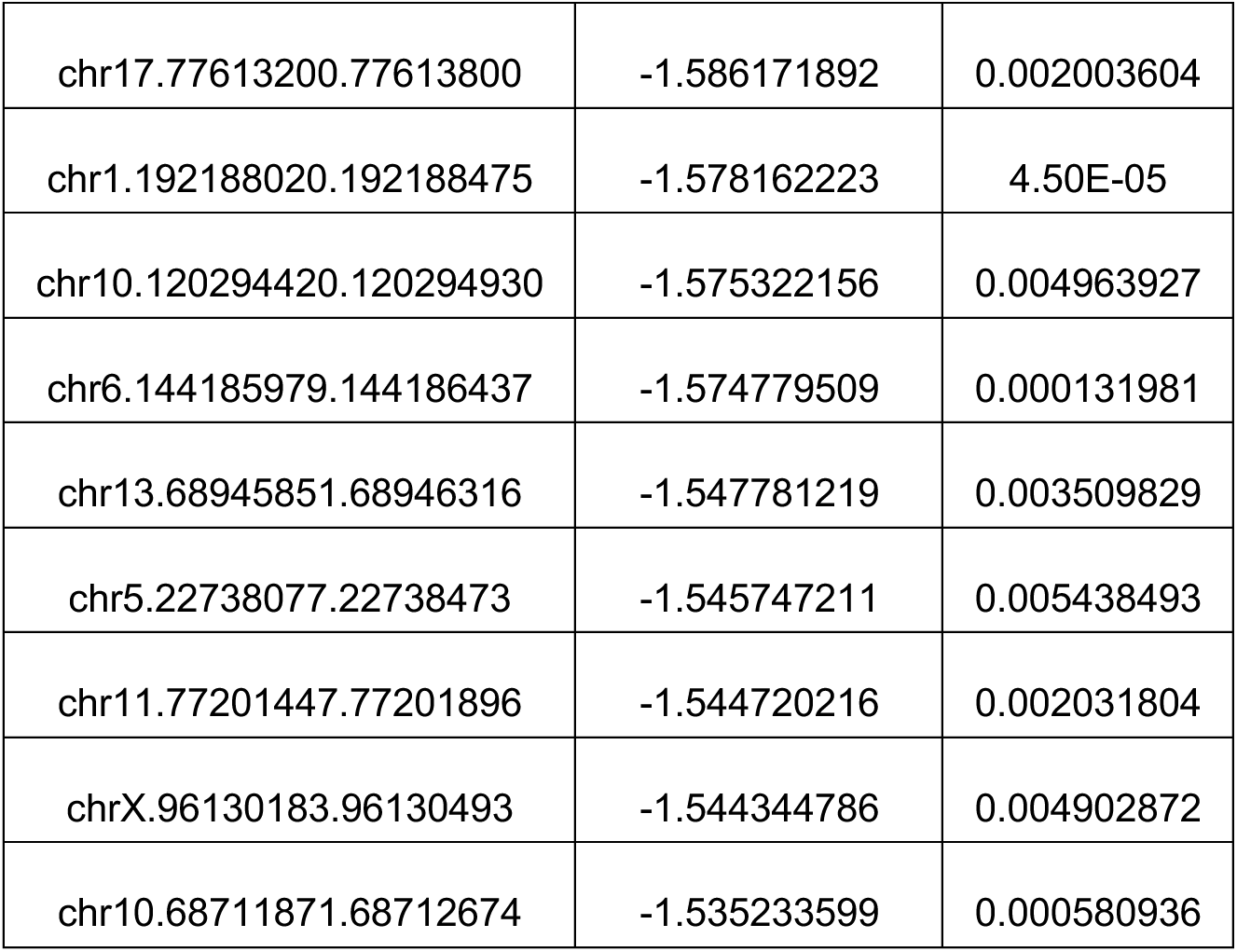
List of top 30 closed peaks in Acvr1^R206H/+^ MEFs compared to Acvr1^+/+^ MEFs.

## Notes

### Competing Interest Statement

The authors have declared no competing interest.

