## Supplemental info for "Hyperactive BMP and Mechanosignaling Remodel Chromatin to Drive Aberrant Osteogenesis in FOP"

^1^Department of Bioengineering, University of Pennsylvania, Philadelphia, PA, 19104, USA, ^2^Department of Orthopaedic Surgery, University of Pennsylvania, Philadelphia, PA, 19104, USA, ^3^Department of Genetics, University of Pennsylvania, Philadelphia, PA, 19104, USA, ^4^The Center for Research in FOP and Related Disorders, University of Pennsylvania, Philadelphia, PA 19104, USA, ^5^Department of Biology, Korea Advanced Institute of Science and Technology, Daejeon, KOR, ^6^Department of Dentistry, College of Dentistry, Kyung Hee University, Seoul, KOR

* Su Chin Heo.

**This PDF file includes:**

Figures S1 to S2

Tables S1 to S4

Figures

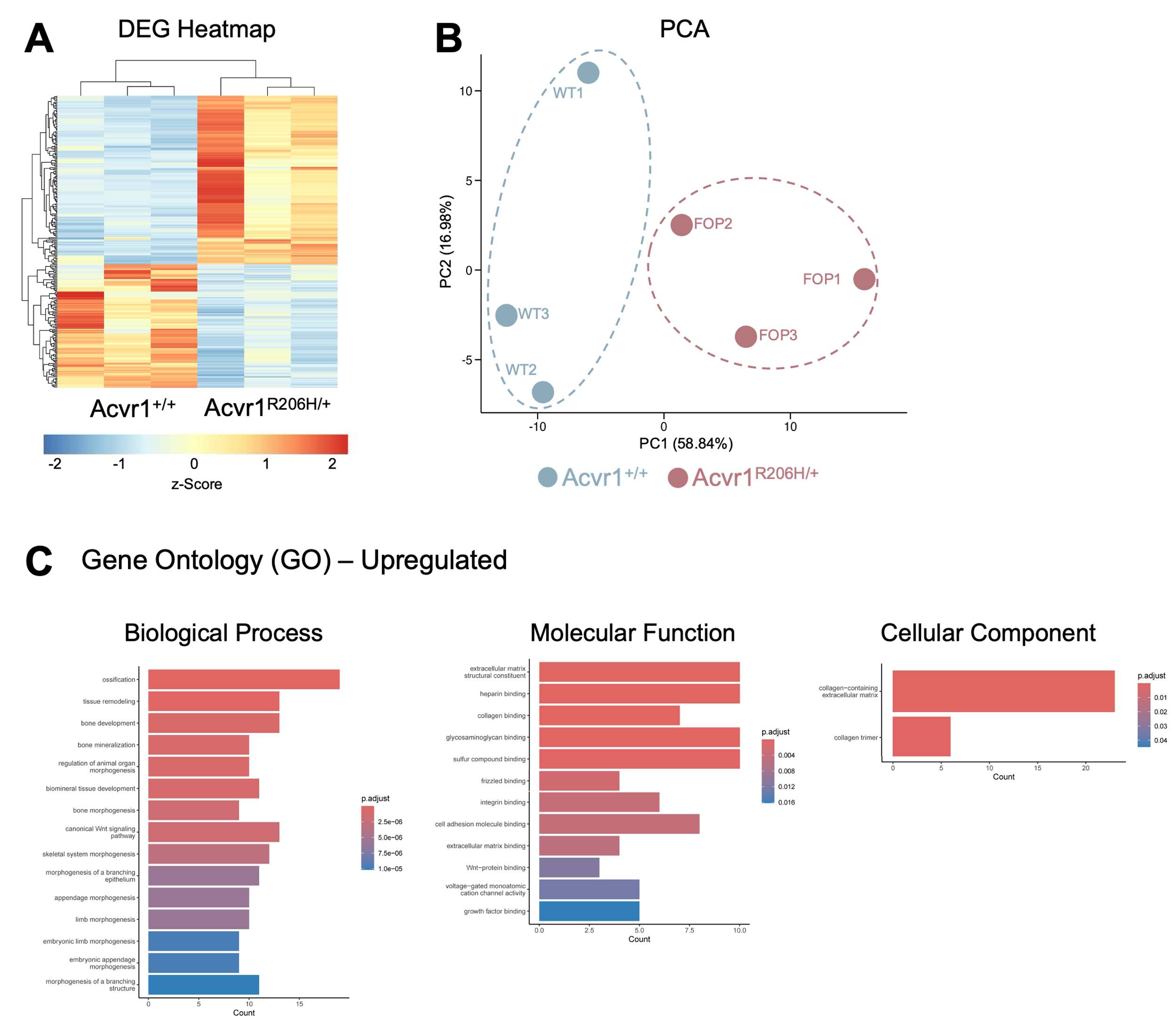

**Figure S1.** **Acvr1^R206H/+^ MEFs display distinct global transcriptomic signatures enriched for osteogenic and matrix-associated pathways. (A)** Heatmap of differentially expressed genes between Acvr1^+/+^ and Acvr1^R206H/+^ MEFs, showing hierarchical clustering of biological replicates (n = 3 biological replicates). **(B)** Principal component analysis (PCA) plot illustrating distinctions between Acvr1^+/+^ and Acvr1^R206H/+^ MEFs based on global gene expression profiles. **(C)** Gene Ontology (GO) enrichment analysis of upregulated genes in Acvr1^R206H/+^ MEFs, highlighting significantly enriched Biological Process, Molecular Function, and Cellular Component terms.

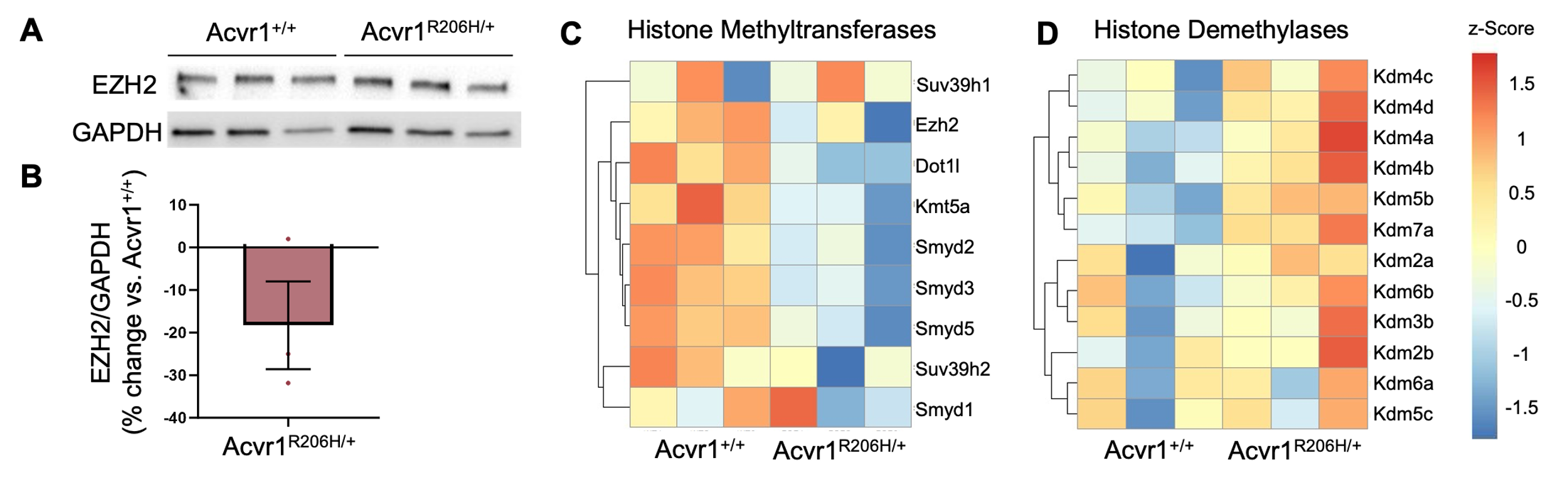

**Figure S2. Altered expression of chromatin-modifying enzymes in Acvr1^R206H/+^ MEFs. (A)** Western blot showing decreased EZH2 protein levels in Acvr1^R206H/+^ MEFs compared to wild-type controls. **(B)** Quantification of relative EZH2 protein expression normalized to GAPDH (n=3 biological replicates). **(C)** Heatmap showing decreased expression of histone methyltransferase genes in Acvr1^R206H/+^ MEFs. **(D)** Heatmap showing increased expression of histone demethylase genes in Acvr1^R206H/+^ MEFs (n=3 biological replicates).

Tables

Table S1. List of top 30 upregulated genes in Acvr1^R206H/+^ MEFs compared to Acvr1^+/+^ MEFs.

| Gene Name | log2FoldChange | padj |
| --- | --- | --- |
| Tnn | 3.479963 | 0.000539 |
| Apol6 | 3.450923 | 2.20E-09 |
| Il20ra | 2.999379 | 0.048819 |
| Icos | 2.827291 | 1.28E-06 |
| Aspn | 2.674363 | 2.04E-05 |
| Fmod | 2.596479 | 4.71E-24 |
| Agt | 2.582337 | 0.008819 |
| Sfrp4 | 2.489648 | 0.000236 |
| Cxcl14 | 2.443428 | 0.001631 |
| Prg4 | 2.438843 | 0.002681 |
| Msx1 | 2.331478 | 2.40E-05 |
| Htr7 | 2.293794 | 0.000446 |
| Kcnk3 | 2.280282 | 0.024489 |
| Prph | 2.270001 | 0.002548 |
| Gm5898 | 2.20729 | 0.005528 |
| Serpinb1a | 2.160948 | 0.002555 |
| Slc26a7 | 2.159098 | 0.004174 |
| Kng2 | 2.150817 | 0.02222 |
| A4galt | 2.136376 | 0.000392 |
| Wnt16 | 2.084207 | 2.44E-06 |
| Abi3bp | 2.057062 | 3.47E-07 |
| Cacna1h | 1.944261 | 5.48E-16 |
| Plxnc1 | 1.931796 | 1.40E-09 |
| Hand1 | 1.906774 | 0.002111 |
| Omd | 1.890677 | 0.000481 |
| Tgfbi | 1.888023 | 2.02E-05 |
| Frmd5 | 1.878517 | 0.000199 |
| Trim30d | 1.859752 | 0.018682 |
| Frem1 | 1.831289 | 0.001585 |
| Ptn | 1.809243 | 3.17E-05 |

Table S2. List of top 30 downregulated genes in Acvr1^R206H/+^ MEFs compared to Acvr1^+/+^ MEFs.

| Gene Name | log2FoldChange | padj |
| --- | --- | --- |
| Matn1 | -4.7162 | 0.003912 |
| Ibsp | -4.28408 | 0.000188 |
| Cyp1a1 | -4.16816 | 1.32E-11 |
| Hsd3b6 | -4.05763 | 4.38E-08 |
| Tie1 | -4.05543 | 0.017371 |
| Slc36a2 | -3.46102 | 0.016654 |
| Krt16 | -3.36207 | 1.71E-05 |
| Cst6 | -3.17748 | 0.000181 |
| F730311O21Rik | -3.1227 | 0.038328 |
| Mal | -3.09639 | 0.000414 |
| Lce1g | -2.99338 | 0.010409 |
| Cnmd | -2.96848 | 4.71E-05 |
| Krt6a | -2.95792 | 1.72E-05 |
| Panx3 | -2.92359 | 0.003899 |
| Sybu | -2.80456 | 0.042238 |
| Tnfsf18 | -2.75598 | 0.031365 |
| Ppp1r1b | -2.75518 | 0.013425 |
| Krt17 | -2.70355 | 4.29E-07 |
| Ppp1r14c | -2.63508 | 0.010044 |
| Prl2c2 | -2.405 | 0.042049 |
| Wnt7a | -2.34222 | 2.38E-05 |
| Prl2c3 | -2.33035 | 0.018017 |
| Epcam | -2.27781 | 0.003619 |
| Cldn3 | -2.27 | 0.000201 |
| Wnt7b | -2.20137 | 0.008529 |
| Apobec2 | -2.19378 | 0.033783 |
| Itgb7 | -2.15829 | 7.83E-08 |
| Ecel1 | -2.15812 | 0.011597 |
| Csn3 | -2.06871 | 0.016662 |
| Trim54 | -2.01693 | 0.01210 |

Table S3. List of top 30 accessible peaks in Acvr1^R206H/+^ MEFs compared to Acvr1^+/+^ MEFs.

| Peaks | log2FoldChange | padj |
| --- | --- | --- |
| chr6.22305950.22306261 | 6.34229268 | 0.006120397 |
| chr16.63600219.63600570 | 5.592941375 | 0.001427996 |
| chr12.36258970.36259257 | 5.381056768 | 0.00182671 |
| chr12.46136355.46136797 | 5.215244666 | 7.19E-06 |
| chr4.98591451.98591748 | 4.961850743 | 0.00043009 |
| chr1.45643494.45643746 | 4.943061668 | 0.00797386 |
| chr1.129893496.129893696 | 4.885326275 | 0.009255883 |
| chr6.53023475.53023801 | 4.73650063 | 0.003115278 |
| chr11.111996407.111997009 | 4.647678569 | 5.76E-09 |
| chr1.179892863.179893108 | 4.585084326 | 0.000937792 |
| chr10.20052232.20052565 | 4.535162954 | 0.000912287 |
| chr7.133155270.133155480 | 4.510613355 | 0.004999723 |
| chr15.56054801.56055142 | 4.419233364 | 4.19E-06 |
| chr16.78069567.78069870 | 4.396055822 | 0.001932931 |
| chrX.56717463.56717761 | 4.381447425 | 0.000232198 |
| chr5.51360306.51360642 | 4.30727699 | 4.35E-06 |
| chr11.63375399.63375604 | 4.28790887 | 0.002223849 |
| chr13.88069857.88070103 | 4.255973257 | 0.003572208 |
| chr2.59552687.59552979 | 4.255973257 | 0.003572208 |
| chr4.98667047.98667423 | 4.253257855 | 9.80E-05 |
| chr18.69639786.69640001 | 4.221743207 | 0.004665329 |
| chr3.146735457.146735689 | 4.207039518 | 0.003348303 |
| chr1.160626782.160627140 | 4.205736592 | 0.001283493 |
| chr7.134707326.134707868 | 4.161083141 | 0.00588362 |
| chr17.48604634.48605039 | 4.12321248 | 0.000171902 |
| chr2.124590483.124590781 | 4.110701203 | 0.008766256 |
| chr9.35436620.35436880 | 4.089638004 | 0.001765702 |
| chr8.115222168.115222368 | 4.084206241 | 0.007140469 |
| chr16.64224428.64224927 | 3.983682461 | 8.72E-06 |
| chr9.70701451.70701748 | 3.922354822 | 0.004486176 |

Table S4. List of top 30 closed peaks in Acvr1^R206H/+^ MEFs compared to Acvr1^+/+^ MEFs.

| Peaks | log2FoldChange | padj |
| --- | --- | --- |
| chr15.36954029.36954318 | -3.981923844 | 5.12E-05 |
| chr15.23317333.23317804 | -2.113734845 | 1.57E-06 |
| chr6.104792308.104792798 | -2.043269651 | 0.000463509 |
| chr4.140419701.140420344 | -1.831806833 | 0.001741442 |
| chr11.77200681.77201195 | -1.821748971 | 1.02E-05 |
| chr5.12936991.12937485 | -1.800962901 | 0.000111101 |
| chr13.43261311.43261778 | -1.768494299 | 0.000148163 |
| chr13.17228379.17228862 | -1.761900489 | 0.007268653 |
| chr4.114960885.114961256 | -1.759477956 | 0.001790703 |
| chr15.43179295.43180025 | -1.741653371 | 1.33E-06 |
| chr5.35967436.35967838 | -1.741104049 | 0.003786315 |
| chr6.144052526.144052956 | -1.736452967 | 0.007903475 |
| chr13.71273549.71273921 | -1.715706797 | 0.007496082 |
| chr13.110403500.110404538 | -1.705842086 | 5.67E-05 |
| chr8.46899031.46899493 | -1.672996855 | 0.000251694 |
| chr2.146610232.146610574 | -1.667723447 | 0.007058031 |
| chr9.14772114.14772459 | -1.626524579 | 0.003663547 |
| chr18.87592491.87593100 | -1.616437121 | 0.004478137 |
| chr8.122013691.122014164 | -1.611909029 | 0.002886263 |
| chr11.66698547.66698946 | -1.607819745 | 0.007382125 |
| chr2.56837252.56838015 | -1.593555598 | 0.00277739 |
| chr17.77613200.77613800 | -1.586171892 | 0.002003604 |
| chr1.192188020.192188475 | -1.578162223 | 4.50E-05 |
| chr10.120294420.120294930 | -1.575322156 | 0.004963927 |
| chr6.144185979.144186437 | -1.574779509 | 0.000131981 |
| chr13.68945851.68946316 | -1.547781219 | 0.003509829 |
| chr5.22738077.22738473 | -1.545747211 | 0.005438493 |
| chr11.77201447.77201896 | -1.544720216 | 0.002031804 |
| chrX.96130183.96130493 | -1.544344786 | 0.004902872 |
| chr10.68711871.68712674 | -1.535233599 | 0.000580936 |
